# Disorganization of basement membrane zone architecture causes impaired melanocyte inhabitation in vitiligo

**DOI:** 10.1101/2022.12.18.520911

**Authors:** Fei Yang, Lingli Yang, Yasutaka Kuroda, Sylvia Lai, Yoshito Takahashi, Tetsuya Sayo, Takeshi Namiki, Kimiko Nakajima, Shigetoshi Sano, Shintaro Inoue, Daisuke Tsuruta, Ichiro Katayama

## Abstract

Vitiligo, a chronic autoimmune skin disorder characterized by selective epidermal melanocyte loss, lacks a well-defined mechanism for this phenomenon. Our study offers compelling insights into vitiligo pathogenesis by revealing disruptions in the basement membrane zone (BMZ) architecture. We observed branched, fragmented, and multilayered lamina densa, accompanied by elevated dermal fibroblast numbers and notable matrix metalloproteinase 2 (MMP2) overexpression. Vitiliginous skin extracts exhibited significant active MMP2 upregulation. To establish a direct link, we intradermally injected MMP2-overexpressing fibroblasts into K14-SCF transgenic mice, resulting in vitiligo-like skin and melanocyte loss, effectively reversed by coadministering MMP2 inhibitors. These groundbreaking findings highlight the pivotal role of disorganized BMZ in vitiligo, proposing MMP2 overexpression in dermal fibroblasts as a potential key contributor. Enhancing our understanding of vitiligo’s mechanisms, this research opens avenues for innovative therapeutic strategies against this challenging autoimmune skin disorder.

**Teaser:** Disrupted skin architecture and MMP2 in dermal fibroblasts hold the key to a potential breakthrough against this puzzling autoimmune disease vitiligo.

## Introductions

Vitiligo, an enigmatic skin disorder with a historical record spanning over 3,500 years, currently impacts 1 to 2% of the global population(*1, 2*), characterized by distinct white patches that can appear localized or disseminated white macules(*3*). Rooted in complexities of genetics, oxidative stress, inflammation, and environmental factors, this acquired, idiopathic, autoimmune cutaneous condition has intrigued researchers for generations. Amidst this intricate interplay, one pivotal question arises: How does the delicate interplay between genetics, environmental stressors, and the immune system give rise to the selective disappearance of epidermal melanocytes, leaving these intriguing white macules in its wake(*4, 5*)? While the role of autoreactive CD8^+^ T cells has been explored in melanocyte destruction(*4, 6*), a deeper exploration into the involvement of other cutaneous cells and the intricacies of the microenvironment surrounding melanocytes in mediating this autoimmune-driven loss continues to elude us.

Under normal homeostatic conditions, epidermal melanocytes, derived from the neural crest, exhibit unique properties, residing in the basal layer of the epidermis in a quiescent state and maintaining a stable ratio with basal keratinocytes(*7*). Perturbations in this delicate balance can lead to intriguing pigmentary disorders(7, 8). The intricate regulation of melanocyte homeostasis involves epidermal keratinocytes, the surrounding stroma, and nearby dermal cells(*9*), with differentiated melanocytes strictly localized at the basement membrane (BM) and reliant on adherence to the basal layers. The BM, a specialized extracellular matrix between the epidermis and dermis(*10, 11*), plays a pivotal role in cellular functions such as adhesion, proliferation, differentiation, and survival(*12*). Despite its well-documented significance in other conditions(*13*) (*14*) (*15*), research on abnormal BM morphology in vitiligo remains limited(*16–19*). Considering the BM’s crucial support for keratinocytes and melanocytes, structural abnormalities within the BM could potentially influence melanocyte disappearance in vitiligo. Central to the constitution of the BM is Collagen IV, the most abundant component, serving as its foundational backbone, complemented by laminin, the BM’s second most abundant component(*20*), synthesized by keratinocytes and fibroblasts. These matrix proteins form a cohesive and sheet-like BM complex(*20*), enabling basal keratinocytes to establish steadfast connections through hemidesmosomes and focal adhesions(*20*).

Matrix metalloproteinases (MMPs), a diverse group of proteolytic enzymes, play a critical role in regulating the cell matrix composition and are essential for various physiological processes(*21*). Dysregulated MMP expression can lead to a wide array of pathological conditions, including inflammation, arthritis, cardiovascular diseases, neurological disorders, and even cancer(*21, 22*). Notably, elevated MMP levels are observed in various autoimmune disorders(*23*) (*24, 25*) (*26*) (*27*) (*28*), indicating their potential involvement in vitiligo.

Building upon this knowledge, our comprehensive investigation, employing electron microscopy, three-dimensional/organotypic culture, and mouse experiments, has unveiled a novel mechanism behind melanocyte disappearance from the BM in vitiligo. The meticulous analysis of lesional skin from vitiligo patients revealed significant aberrations in the BM’s structure and components compared to healthy skin, with fibroblast mediated MMP2 overexpression implicated as a key factor leading to BM matrix degradation and subsequent melanocyte detachment. Our findings mark a meaningful milestone in vitiligo research, offering deeper insights into the disease’s etiology and pathogenesis.

## Results

### Basement membrane zone architecture is disorganized in vitiligo

To investigate the BM zone (BMZ) in vitiligo, we performed an ultrastructural analysis using biopsy specimens from both healthy donors (n = 5) and patients with generalized vitiligo (two stable and three progressive cases). We observed abnormal BM morphology in samples from both stable and progressive vitiligo patients. Specifically, healthy skin displayed intact lamina densa with high electron density, while vitiliginous skin showed branched and fragmented lamina densa, some of which were multilayered. the BM of healthy skin contained intact lamina densa with high electron density, whereas the BM of vitiliginous skin contained branched and fragmented lamina densa, some of which were multilayered (**Fig 1a**). Moreover, the anchoring fibrils, which were normally distributed in healthy skin, were scattered and distributed in both the upper and lower portions of the branched lamina densa in vitiliginous skin (**Fig 1a, red asterisks**).

**Figure 1.**
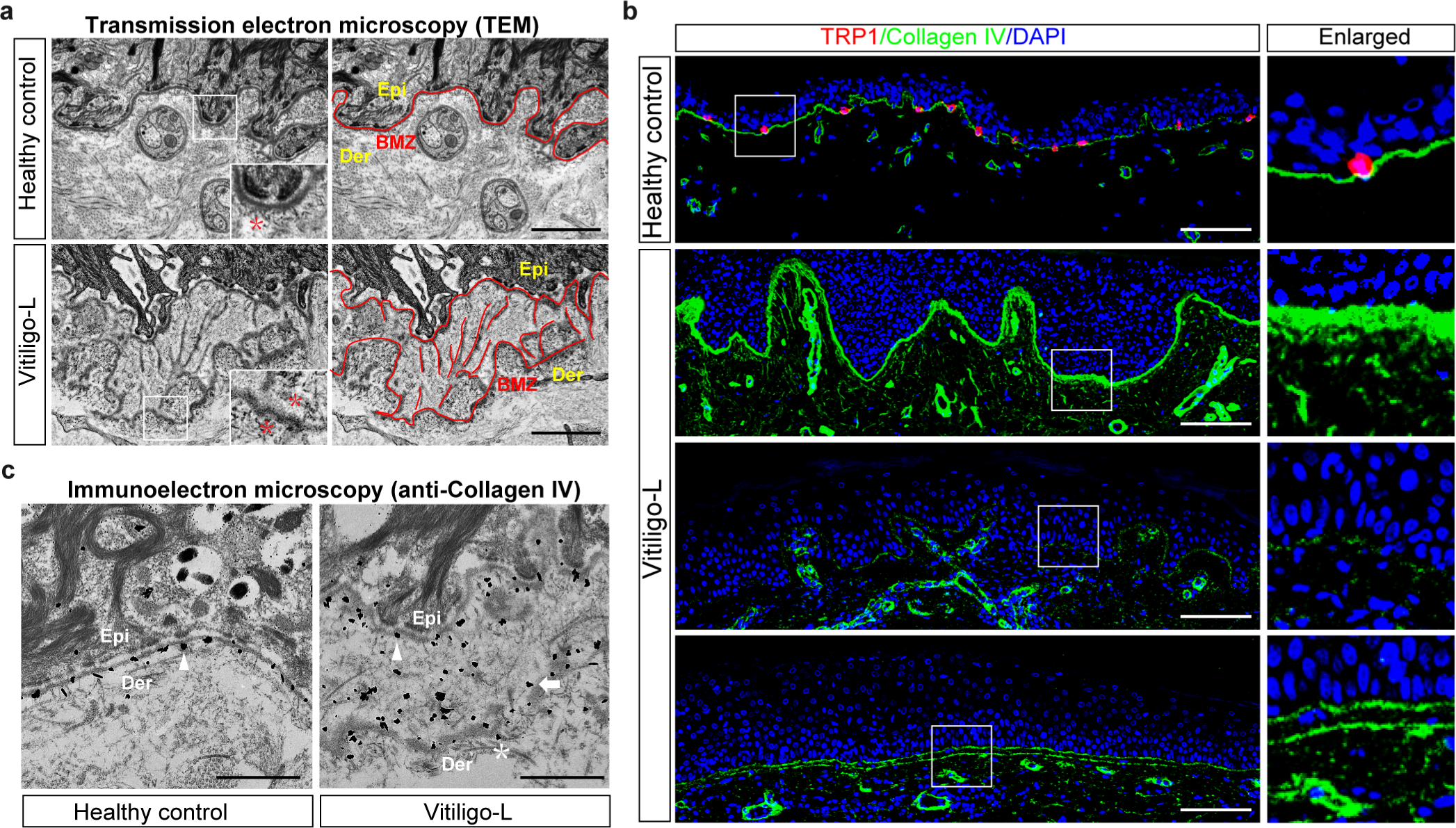
Disorganization of basement membrane zone architecture in vitiligo **a,** Representative ultrastructural images of ultrathin sections of skin from five healthy controls and five patients with generalized vitiligo. In the right panel, the areas of epidermis and dermis are represented by Epi and Der, respectively, and the basement membrane zone is indicated by a red line. In the left panel, the area of the white rectangle is shown enlarged at the lower right. Red asterisk: anchoring fibril; Epi: epidermis; Der: dermis; BMZ: basement membrane zone. Scale bars: 2 μm. **b,** Representative immunofluorescence images of skin specimens from five healthy controls and 11 patients with generalized vitiligo stained with anti-collagen IV (green), anti-TRP1 (red), and DAPI (blue). The areas in the white rectangles are shown in the corresponding right panels. Scale bar: 100 μm. **c,** Representative immunoelectron microscopy images of frozen sections stained with anti-collagen IV. White arrowhead: collagen IV-labeled gold-particle deposits close to the lamina densa; White arrow: collagen IV-labeled gold-particle deposits close to the branched structure of lamina densa; White asterisk: collagen IV-labeled gold-particle deposits close to the layered structure of lamina densa. Scale bars: 1 μm.

Next, we used immunofluorescence staining to investigate the expression and localization of BM proteins. The BM in healthy skin exhibited normal morphology with a thin continuous linear band of collagen IV and a normal number and distribution of melanocytes. In contrast, vitiliginous skin displayed various abnormal BM morphologies, including obvious thickening, severe disruption, and multilayering, accompanied by the disappearance of melanocytes (**Fig 1b**).

We further investigated the ultrastructural localization of collagen IV using immunoelectron microscopy. In healthy skin, we observed intact BM with collagen IV-labeled gold-particle (**Fig 1c, white arrowhead in left panel**) deposits located close to the lamina densa in a continuous arrangement. However, in vitiliginous skin, we observed conspicuous structural alterations of the BM, with collagen IV-labeled gold-particle deposits scattered across a wide area not only close to the normal lamina densa area (**Fig 1c, white arrowhead in right panel**) but also near the branched (**Fig 1c, white arrow**) and multilayered structures (**Fig 1c, white asterisk**) of lamina densa. These results revealed a disrupted BMZ architecture in lesional vitiliginous skin.

### Matrix metalloproteinase 2 expression is elevated in the dermis of vitiliginous skin

To investigate the cause of BM structure disruption, we measured the expression levels of MMP2 and MMP9, the major proteolytic enzymes capable of degrading BM collagen, in skin sections from vitiligo patients. We found that vitiliginous skin displayed significantly elevated MMP2 expression in the upper dermis (**Fig 2a and 2b**).

**Figure 2.**
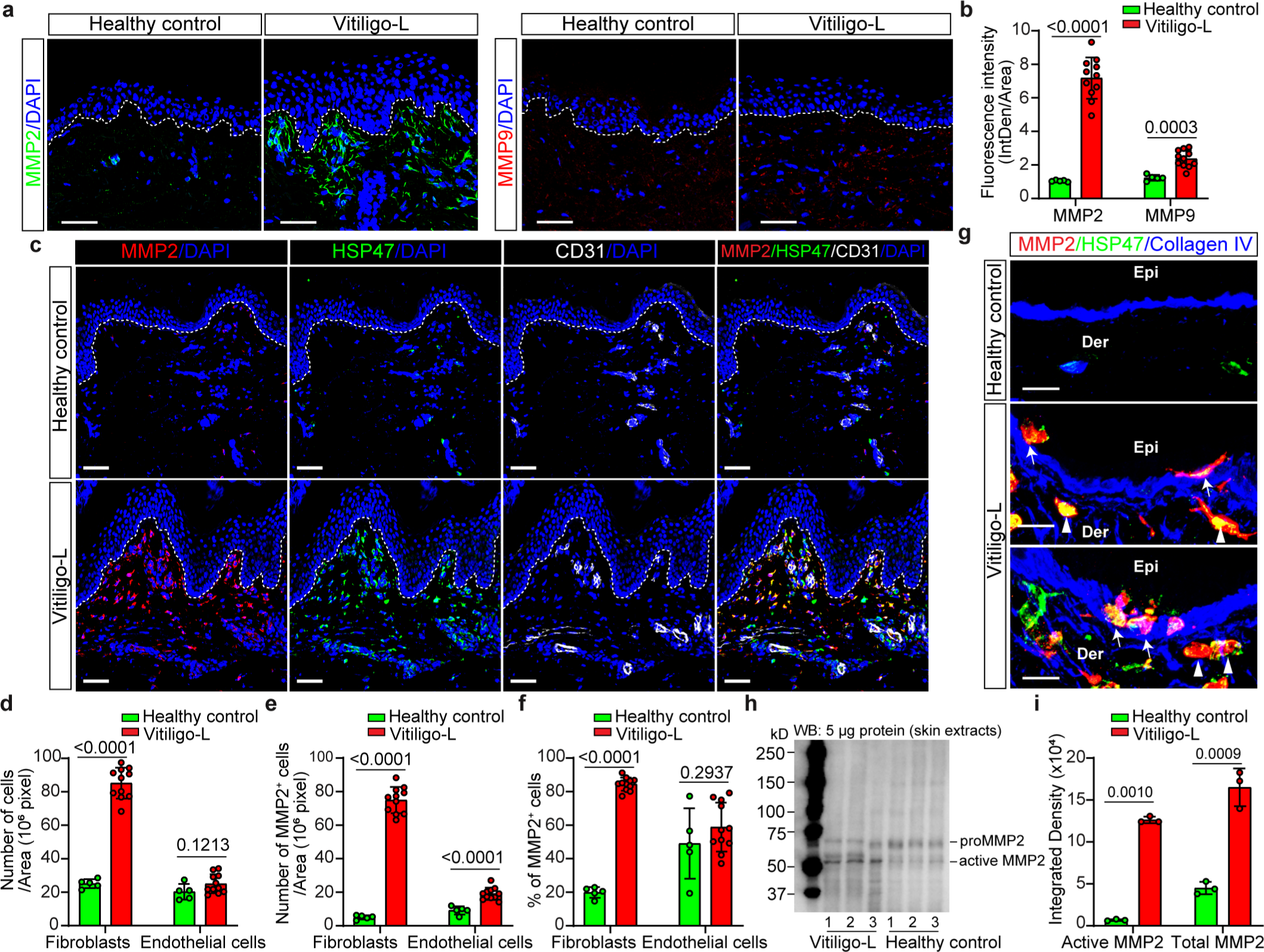
Matrix metalloproteinase 2 expression is elevated in the dermis of vitiliginous skin **a,** Representative immunofluorescence images of frozen sections of skin from five healthy controls and 11 patients with generalized vitiligo. MMP2 is stained green, and MMP9 is stained red. Vitiligo- L: lesional vitiliginous skin. **b,** Quantitation of MMP2 and MMP9 expression by fluorescence intensity measurement. **c,** Representative immunofluorescence staining of MMP2, HSP47, and CD31 in skin from healthy controls and patients with vitiligo. **d,** Quantitation of fibroblast and endothelial cell numbers in healthy and vitiliginous skin. **e,** Quantitation of MMP2^+^ fibroblast and MMP2^+^ endothelial cell numbers. **f,** The ratios of MMP2^+^ fibroblasts among all fibroblasts and MMP2^+^ endothelial cells among all endothelial cells. **g,** Representative immunofluorescence staining of MMP2^+^ fibroblasts distributed around the BM. MMP2 is stained red; the fibroblast marker HSP47 is stained green; and collagen IV, the main component of BM, is stained blue. White arrowhead: MMP2^+^ fibroblasts close to the BM; White arrow: MMP2^+^ fibroblasts passed through the BM. **h,** Western blot analyses showing expression of the pro and active forms of MMP2 in skin lysates from healthy skin and lesional vitiliginous skin. **i,** Quantitation of active MMP2 and total MMP2 (the total of active and pro MMP2) expression according to western blot data. Nuclei are counterstained in blue with DAPI in **a** and **c**. Scale bars in **a** and **c**: 50 μm. Scale bars in **g**: 20 μm.

Fibroblasts(*29*) and vascular endothelial cells(*30*) are the primary sources of MMP2 production in human skin. To identify the main contributors to elevated MMP2 levels in vitiliginous skin, we conducted immunolocalization studies on healthy and vitiligo-affected skin tissues. Frozen sections from five healthy controls and 11 vitiligo patients were stained with anti-MMP2 antibodies, along with anti-HSP47 (fibroblast marker) and anti-CD31 (endothelial cell marker) antibodies.

In the vitiliginous skin, fibroblast numbers were significantly higher compared to healthy skin, while endothelial cell numbers showed no significant difference (**Fig 2c and 2d**). Moreover, MMP2-positive fibroblasts in vitiliginous skin were nearly four times more abundant than in healthy skin, with no significant difference in MMP2-positive endothelial cells (**Fig 2e and 2f**).

To investigate MMP2-positive fibroblasts’ localization in relation to the BM, we conducted immunofluorescent labeling using anti-MMP2 antibodies, and co-stained with anti-HSP47 and anti-collagen IV (a BM marker) antibodies. Interestingly, MMP2-positive fibroblasts in vitiliginous skin were found in proximity to the BM (**white arrowheads in Fig 2g**) and were observed traversing the BM to reach the epidermis (**white arrows in Fig 2g**).

Next, we examined the activity of MMP2 by western blot using skin lysates from three patients with vitiligo (one stable, two progressive) and three healthy controls. MMP2 was highly expressed and mainly in its active form in vitiliginous skin, in contrast to healthy skin where it was slightly expressed and mainly in the inactive form (**Fig 2h and 2i**). These findings indicated that increased MMP2 expression by dermal fibroblasts in vitiliginous skin may contribute to BM destruction and subsequently, melanocyte disappearance.

### Overexpression of MMP2 in dermal fibroblasts results in melanocyte disappearance

To determine whether damage to the BM precedes melanocyte disappearance, we developed an *in vitro* 3D skin equivalent model. In this model, half of the collagen gel in the dermis layer was prepared with mock-transfected fibroblasts (Mock-Fbs) and half with MMP2-overexpressing fibroblasts (MMP2-Fbs) (**Fig 3a**). Once the structured collagen gel was established, melanocytes and keratinocytes were uniformly seeded on top of the gel. After 1 week of air-lift culture, macroscopically pigmented skin developed on the part of the gel prepared with Mock-Fbs, whereas no pigmentation developed on the part of the gel prepared with MMP2-Fbs (**Fig 3b**). The 3D skin equivalents prepared with MMP2-Fbs displayed damaged BM and a lack of melanocytes, while those prepared with Mock- Fbs had a typical BM morphology and distributed melanocytes in the basal layer (**Fig 3c**).

**Figure 3.**
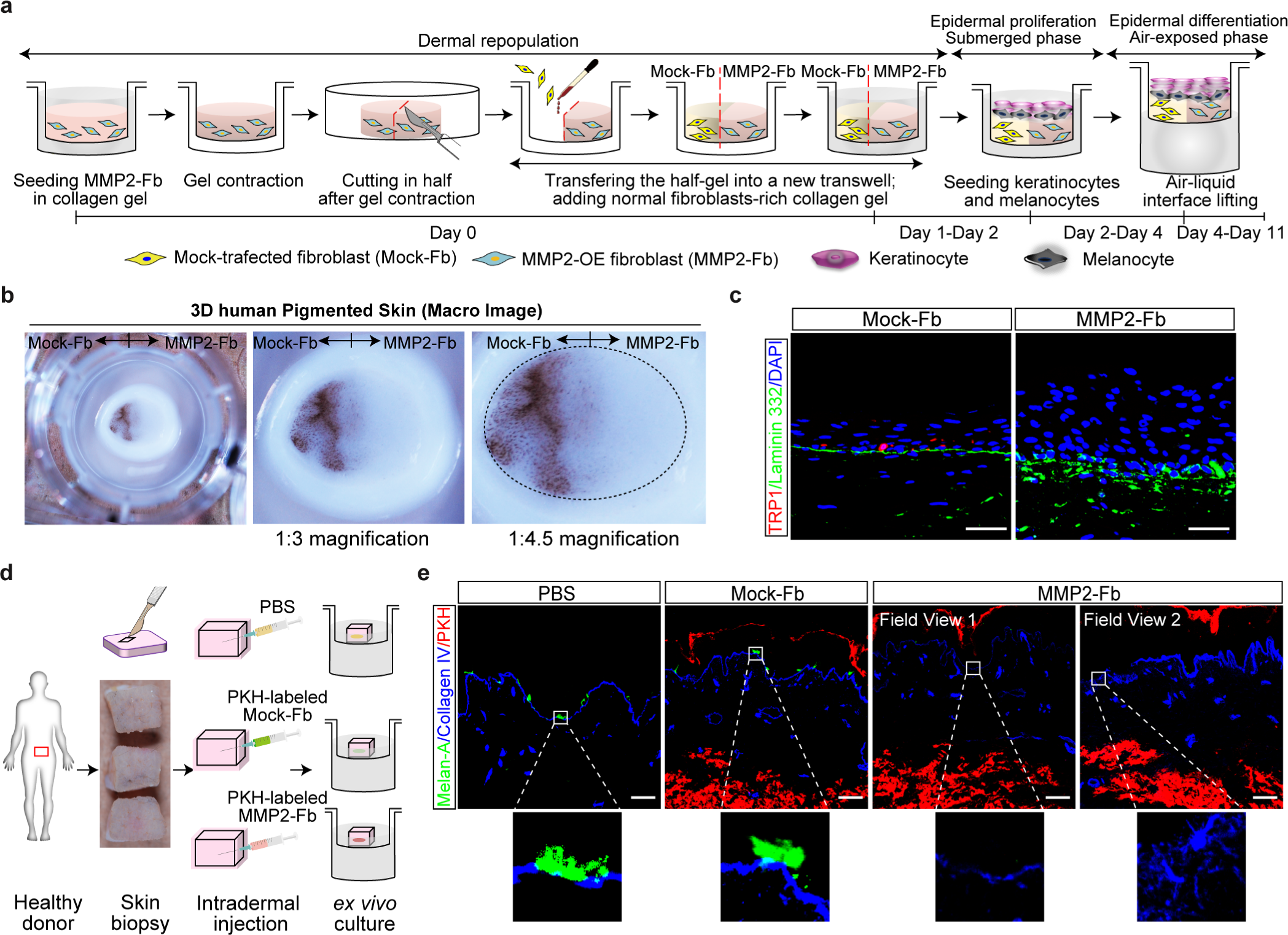
Overexpression of MMP2 in dermal fibroblasts results in melanocyte disappearance in three-dimensional human full-thickness skin equivalent and human *ex vivo* cultured skin explants. **a,** Schematic flow diagram depicting the reconstructed three-dimensional human skin full-thickness equivalent model. **b,** Macroscopic images of skin equivalents. **c,** Representative immunofluorescence staining of the melanocyte marker TRP1 and the BM marker laminin 332 in skin equivalents. TRP1 is stained red, and laminin 332 is stained green. Nuclei are counterstained in blue with DAPI. **d,** Schematic diagram of the human skin explant culture model. **e,** Representative immunofluorescence staining of collagen IV and melan-A in cultured human skin explants. Collagen IV is stained blue, and melan-A is stained green. Intradermally injected fibroblasts are labeled with PKH fluorescent dye in red. The white rectangle outlines the area of melanocytes and BM, and the detail is shown in the corresponding lower panel. Scale bars in **c** and **e**: 50 μm. Data in **b**, **d**, **e**, **f,** and **i** are shown as mean ± standard deviation.

We further created an *ex vivo* subcutaneous injection model, in which Mock-Fbs or MMP2-Fbs were subcutaneously inoculated into cultured, full-thickness, fresh skin explants. To track the injected fibroblasts, we labeled them with PKH fluorescence cell-membrane labeling dye (**Fig 3d**). Skin explants injected with MMP2-overexpressing fibroblasts displayed BM disruption and melanocyte disappearance (**Fig 3e**). These results indicated that increased MMP2 expression by dermal fibroblasts is a key factor in melanocyte disappearance in vitiligo.

### Melanocyte disappearance results from decreased adhesion to the BM

We performed immunofluorescent staining to further examine the disappearance of melanocytes from the damaged BM. Immunofluorescent staining revealed that melanocytes in depigmented 3D skin equivalents tended to be located at the suprabasal layer, detached from the BM, and were negative for cleaved-caspase 3, suggesting that apoptotic events were not involved in their disappearance (**Fig 4a**).

**Figure 4.**
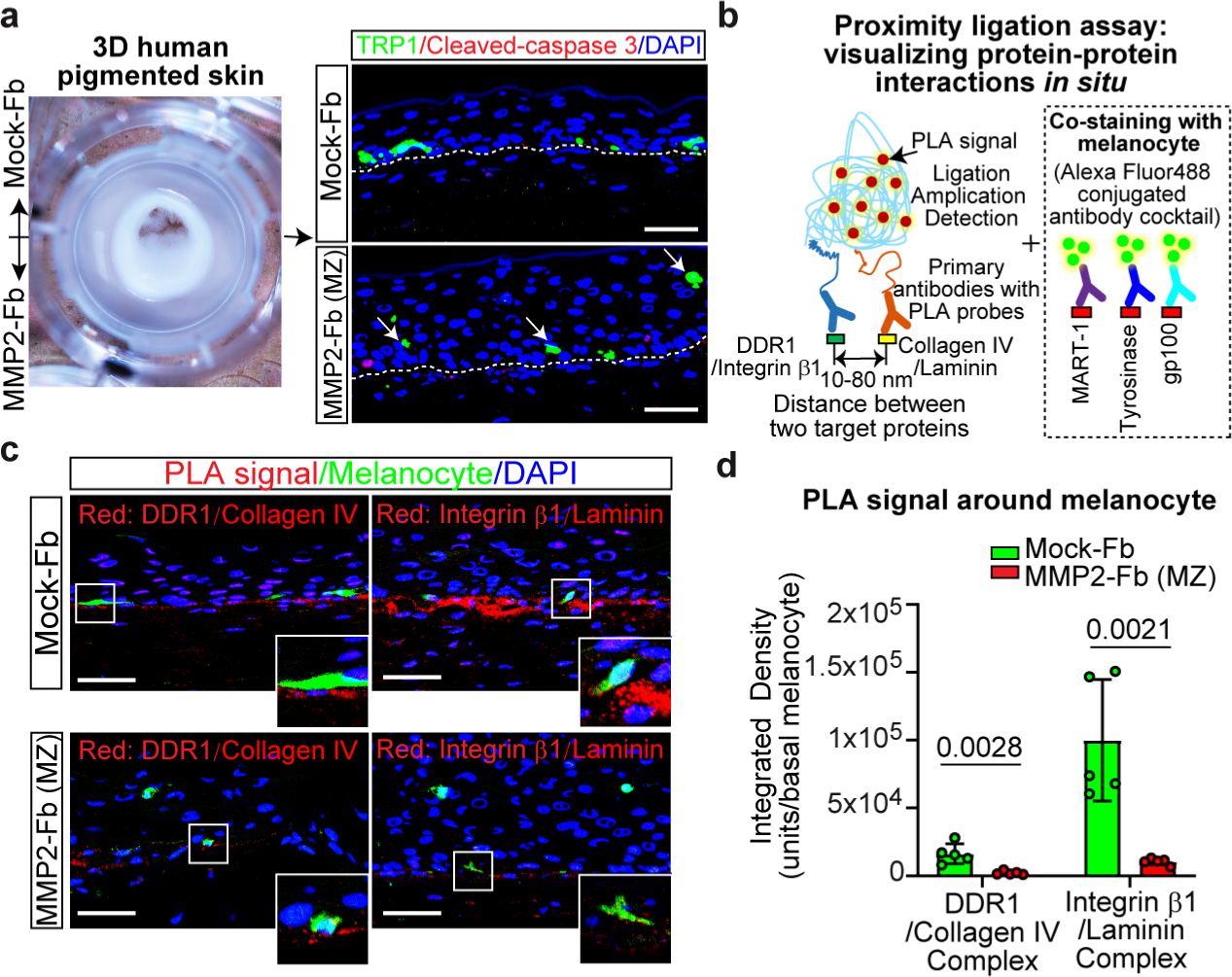
Melanocyte disappearance results from decreased adhesion to the BM in the three-dimensional skin equivalent model. **a,** Representative immunofluorescence staining of TRP1 and cleaved-caspase 3 in three-dimensional skin equivalents. TRP1 is stained green, and cleaved-caspase 3 is stained red. Mock-Fb indicates skin equivalent embedded with mock-transfected fibroblasts; MMP2-Fb (MZ) indicates the marginal zone of skin equivalent embedded with MMP2-overexpressing fibroblasts. Detached melanocytes are shown by white arrows. **b,** Schematic diagram of the proximity ligation assay and melanocyte antibody cocktail immunofluorescence co-staining. **c,** The interactions between DDR1 and collagen IV and between integrin β1 and laminin in skin equivalents are visualized in red fluorescence by proximity ligation assay. Melanocytes are visualized in green. Nuclei are counterstained in blue with DAPI. The areas in the white rectangles are shown at the lower right. **d,** Quantitation of the fluorescence intensity around melanocytes in proximity ligation assays of five skin equivalents. Scale bars in **a** and **c**: 50 μm. Data in **d** are shown as mean ± standard deviation.

Cells in the basal layer of the epidermis are bound to the BM mainly through integrins, but the mechanistic details of melanocyte adhesion to the BM are still unknown. Melanocytes have been reported to express integrin β1 both *in vitro* and *in vivo*(*31*). In addition, discoidin domain receptor 1 (DDR1), a membrane protein, was recently reported to bind collagen IV and mediate the adhesion of melanocytes to BM(*32*). Furthermore, some studies reported that loss of melanocytes in vitiligo resulted from decreased DDR1 expression in melanocytes(*33, 34*). We examined adhesion patterns between melanocytes and the BM matrix using *in situ* proximity ligation assay (PLA), an antibody-based technology that enables the visualization of protein–protein interactions. Specifically, we examined integrin β1–laminin-binding and DDR1–collagen IV binding in 3D skin equivalents. In the PLA, when a pair of specific primary antibodies (DDR1 and collagen IV; integrin β1 and laminin) is in close proximity, complementary DNA strands on a corresponding pair of secondary antibodies with PLA probes engage in rolling circle amplification and generate a single fluorescent signal *in situ*, indicating the presence of the corresponding protein–protein interaction (**Fig 4b**). This assay revealed decreased interactions of DDR1–collagen IV and integrin β1–laminin in 3D skin equivalents prepared with MMP2-overexpressing fibroblasts (**Fig 4c and 4d**). Abnormal adhesion between melanocytes and the BM was implicated in their disappearance.

Immunofluorescence staining detected melanocyte detachment from the BM in perilesional skin of vitiligo patients (**Fig 5a**). Expression levels of DDR1, collagen IV, integrin β1, and laminin showed no significant difference between healthy and perilesional vitiliginous skin (**Fig 5b, 5c, and 5f**). We employed *in situ* PLA and co-immunohistochemistry staining of melanocytes with anti-TRP1 antibody to visualize adhesions between melanocytes and the BM matrix (**Fig 5d and 5e, white rectangle box in lower right panel**). In healthy skin melanocytes, integrin β1–laminin interactions were more predominant compared to DDR1–collagen IV interactions. However, both interactions were significantly diminished in perilesional vitiliginous skin melanocytes (**Fig 5e and 5g**). These findings suggest that melanocyte loss in vitiligo may be linked to a reduced adhesion pattern between melanocytes and the BM (**Fig 5h**).

**Figure 5.**
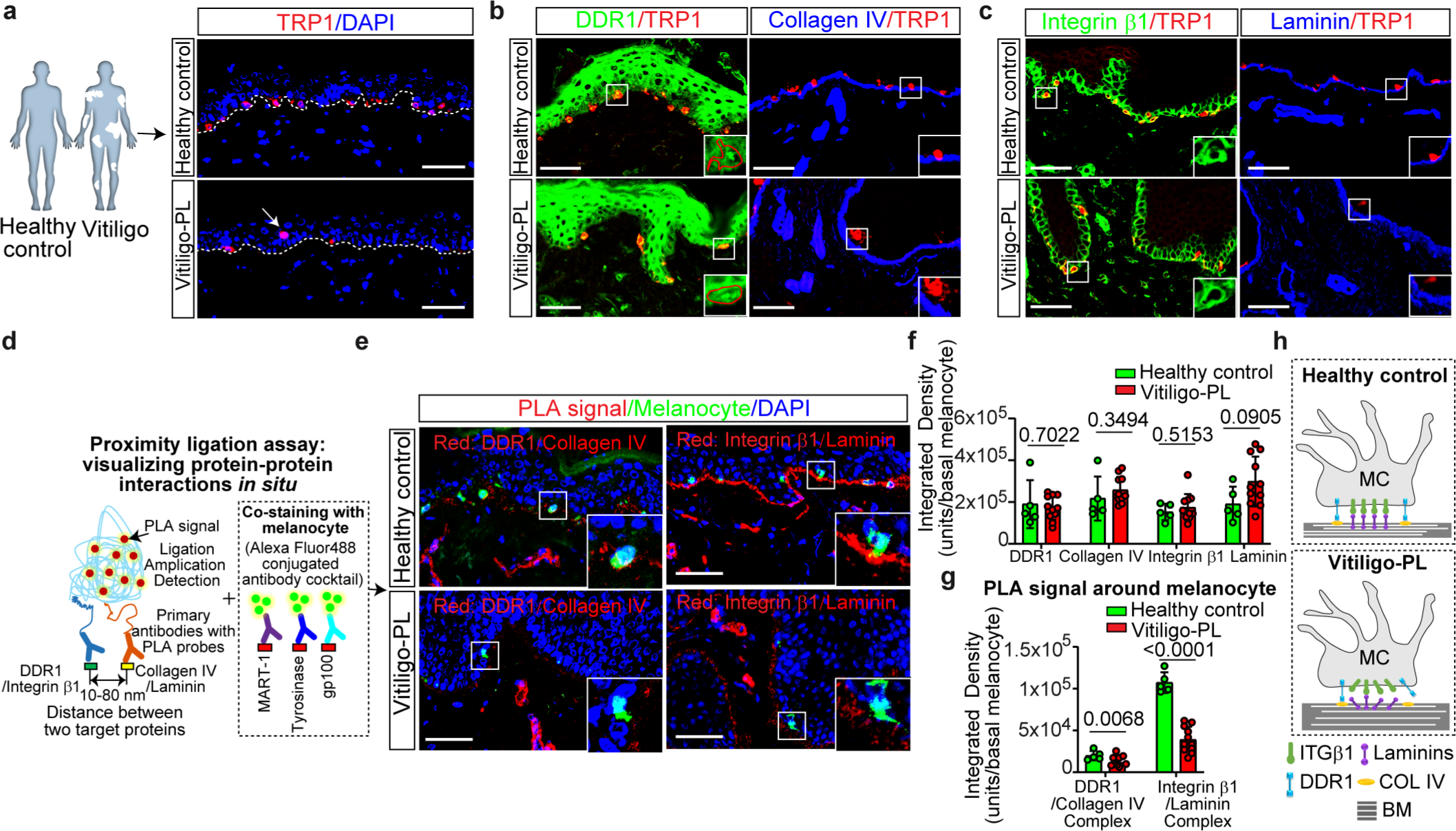
Melanocyte disappearance results from decreased adhesion to the BM in vitiliginous skin. **a,** Representative immunofluorescence staining of TRP1 in healthy skin and perilesional vitiliginous skin. TRP1 is stained red. Nuclei are counterstained in blue with DAPI. Detached melanocytes are shown by white arrows. Vitiligo-PL indicates perilesional vitiliginous skin. **b,** Representative immunofluorescence staining of DDR1, collagen IV, and TRP1 in healthy skin and perilesional vitiliginous skin. DDR1 is stained green, collagen IV is stained blue, and TRP1 is stained red. The area in the white rectangle is shown on the lower right. The red lines circling the areas in the enlarged images indicate melanocytes. **c,** Representative immunofluorescence staining of integrin β1, laminin, and TRP1 in healthy skin and perilesional vitiliginous skin. Integrin β1 is stained green, laminin is stained blue, and TRP1 is stained red. The area in the white rectangle is shown on the lower right. **d,** Schematic diagram of the proximity ligation assay and melanocyte antibody cocktail immunofluorescence co-staining. **e,** The interactions between DDR1 and collagen IV and between integrin β1 and laminin are visualized in red fluorescence by proximity ligation assay. Melanocytes are visualized in green. Nuclei are counterstained in blue with DAPI. The area in the white rectangle is shown on the lower right. **f,** Quantitation of the immunofluorescence signal around melanocytes in **b** and **c**. **g,** Quantitation of the fluorescence intensity around melanocytes in proximity ligation assays in **e**. **h,** Illustration of melanocyte–BM adhesion in healthy skin and vitiliginous skin. Scale bars in **a, b, c,** and **e**: 50 μm. Data in **f** and **g** are shown as mean ± standard deviation.

Unlike melanocyte detachment, keratinocyte detachment (blistering) was not observed in depigmented 3D skin equivalents and perilesional vitiliginous skin, despite weakened focal adhesion signal (integrin β1–laminin binding) around basal keratinocytes (**Fig 4c**; **Fig 5e**). Hence, we investigated hemidesmosome protein expression and binding signals. Expression levels of integrin α6 and laminin showed no difference between healthy and lesional vitiliginous skin **(Fig 6a and 6c)**. However, collagen XVII and collagen IV levels were significantly elevated in the lesional vitiliginous skin (**6b and 6c**). Using *in situ* PLA, we examined hemidesmosome adhesion signals between basal keratinocytes and the BM matrix (**Fig 6d**). In healthy skin basal keratinocytes, integrin α6–laminin interactions predominated over collagen XVII–collagen IV interactions (**Fig 6e and 6f**). In contrast, basal keratinocytes of vitiliginous skin exhibited a marked reduction in integrin α6–laminin interaction, while collagen XVII–collagen IV interaction notably increased (**Fig 6e and 6f**). These results suggest altered adhesion patterns between basal keratinocytes and the BM (**Fig 6g**). Compensatory enhanced hemidesmosome adhesion, possibly facilitated by collagen XVII binding to collagen IV, may underlie the absence of skin blistering.

**Figure 6.**
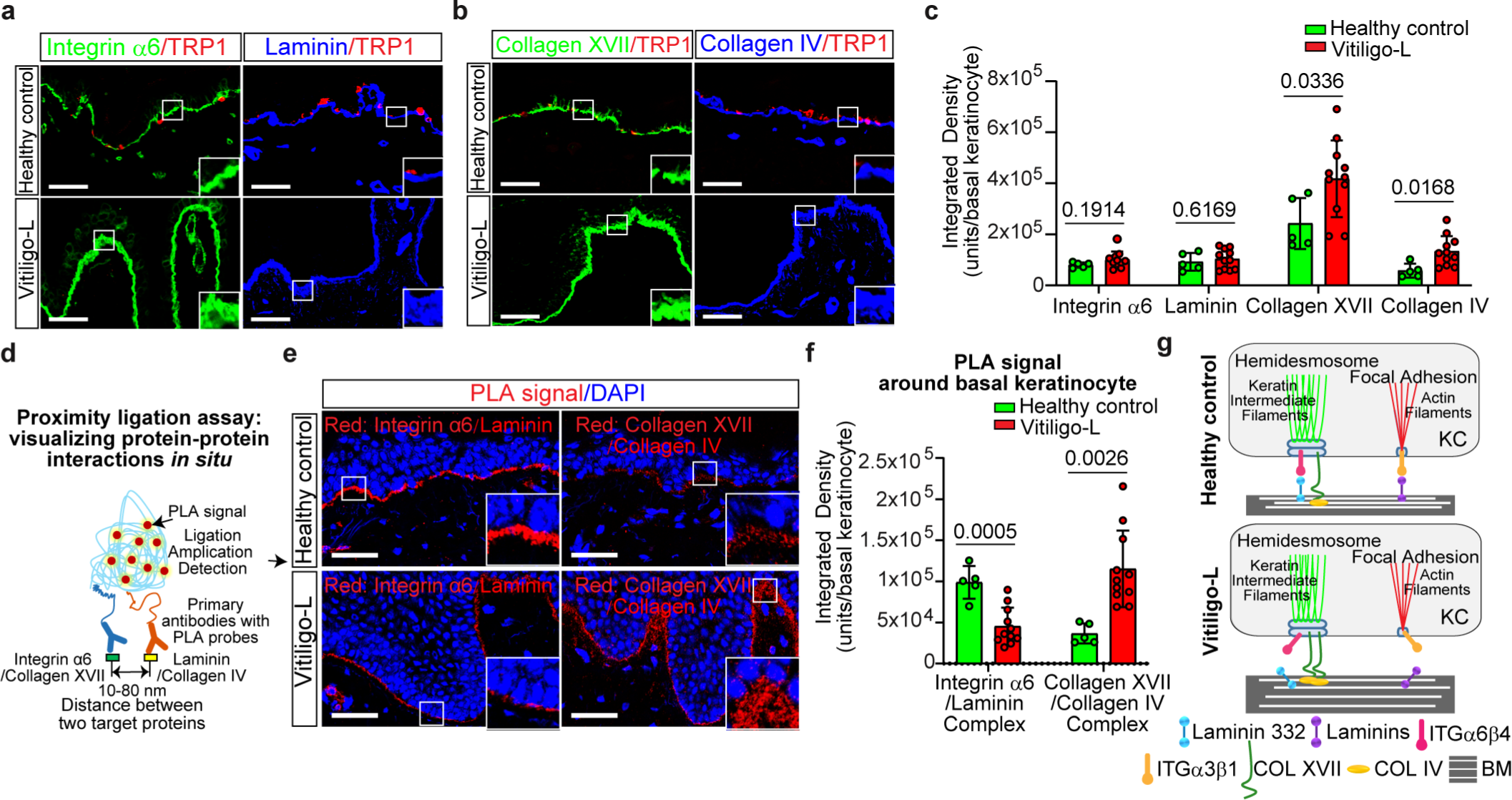
Compensatory enhanced hemidesmosome adhesion in the basal keratinocytes of vitiliginous skin. **a,** Representative immunofluorescence staining of integrin α6, laminin, and TRP1 in healthy skin and lesional vitiliginous skin. Integrin α6 is stained green, laminin is stained blue, and TRP1 is stained red. The area in the white rectangle is shown on the lower right. **b,** Representative immunofluorescence staining of collagen XVII, collagen IV, and TRP1 in healthy skin and lesional vitiliginous skin. Collagen XVII is stained green, collagen IV is stained blue, and TRP1 is stained red. The area in the white rectangle is shown on the lower right. **c,** Quantitation of the immunofluorescence signal around basal keratinocytes in **a** and **b**. **d,** Schematic diagram of the proximity ligation assay. **e,** The interactions between integrin α6 and laminin and between collagen XVII and collagen IV are visualized in red fluorescence by proximity ligation assay. Nuclei are counterstained in blue with DAPI. The area in the white rectangle is shown on the lower right. **f,** Quantitation of the fluorescence intensity around basal keratinocytes in the proximity ligation assays in **e**. **g,** Illustration of basal keratinocyte–BM adhesion in healthy skin and vitiliginous skin. Scale bars in **a, b,** and **e**: 50 μm. Data in **c** and **f** are shown as mean ± standard deviation.

### Decreased melanocyte adhesion to the BM is caused by abnormal BM architecture

The expression levels of DDR1, collagen IV, integrin β1, and laminin were similar between vitiliginous and healthy skin. However, vitiliginous skin exhibited decreased DDR1–collagen IV and integrin β1–laminin interactions (**Fig 5**). To investigate the cause of these reduced interactions, we assessed the physical distance between melanocytes and the BM using electron microscopy (**Fig 7a**). As basal cells typically attach to the lamina densa of the BM, we measured the width of the lamina lucida (the distance between the basal cell plasma membrane and the lamina densa) in both healthy and vitiliginous skin. Surprisingly, we found no difference between the two (**Fig 7a and 7c**).

**Figure 7.**
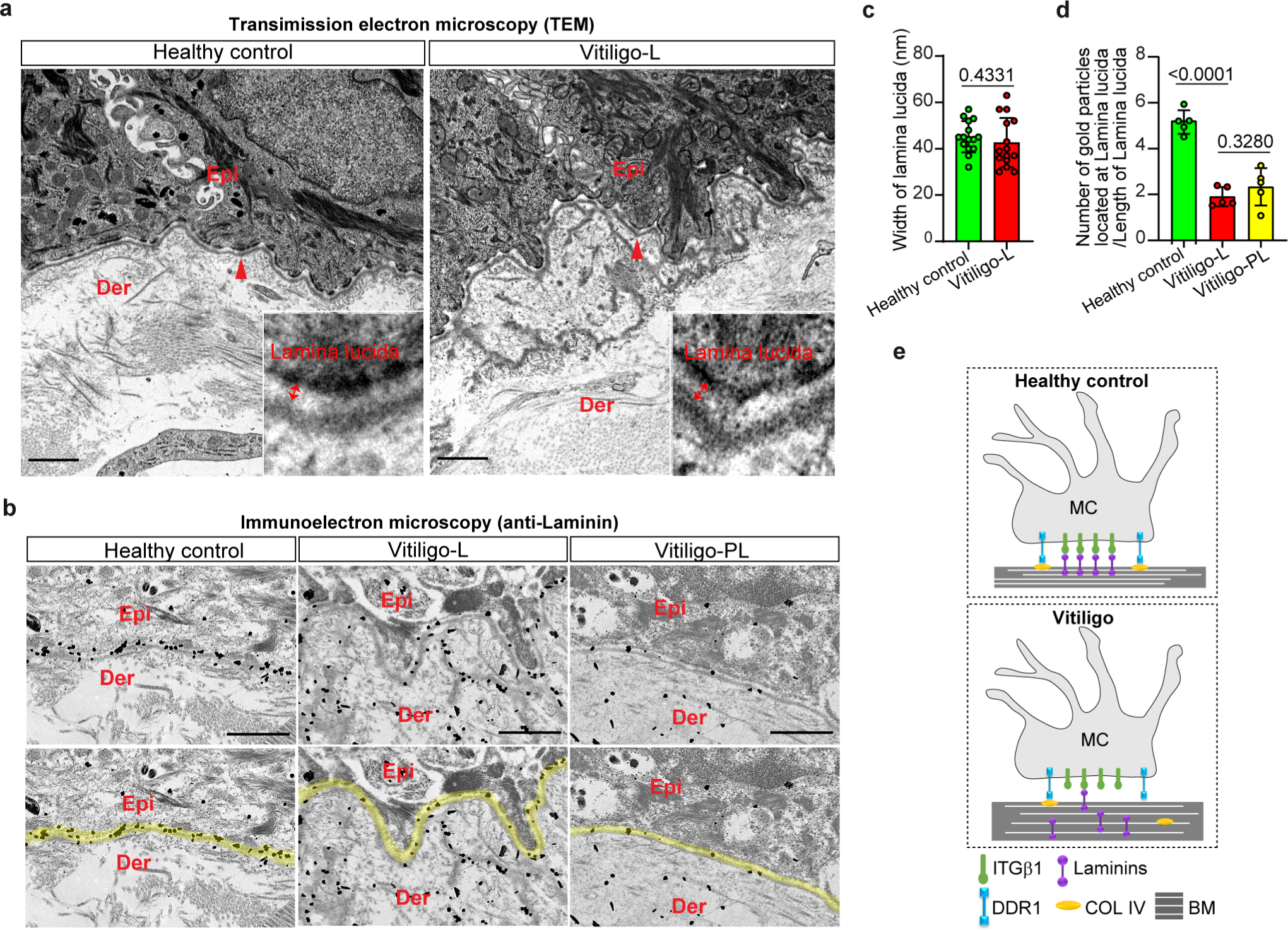
The decreased adhesion of melanocytes to the BM is due to abnormal BM architecture. **a,** Representative ultrastructural images of ultrathin sections of skin from five healthy controls and five patients with generalized vitiligo. An enlarged image of the area indicated by the red arrowhead is shown on the lower right. The red double arrow indicates the lamina lucida. Epi: epidermis; Der: dermis. Scale bars: 1 μm. **b,** Representative immunoelectron microscopy images of ultrathin skin sections stained with anti-laminin antibody. The area of the lamina lucida is outlined in yellow. Scale bars: 1 μm. **c,** Quantitation of the width of the lamina lucida in **a**. Three measurements were performed per section. **d,** Quantitation of the ratio of the number of laminin gold particles located at the lamina lucida to the length of the lamina lucida. **e,** Illustration of melanocyte–BM adhesion in healthy skin and vitiliginous skin. Data in **c** and **d** are shown as mean ± standard deviation.

To further explore the decreased binding, we performed immunoelectron microscopy. In healthy skin, laminin, the primary adhesion molecule responsible for melanocyte binding to the BM, was clearly distributed along and confined to the lamina densa. However, in lesional and perilesional vitiliginous skin, laminin was diffusely spread within the lamina densa and beneath the disorganized BM architecture (**Fig 7b**). Moreover, the amount of laminin confined in the lamina lucida was significantly reduced in vitiliginous skin compared to healthy skin (**Fig 7b and 7d**). These findings suggest that the disorganized arrangement of the basement membrane zone (BMZ) may contribute to the loss of melanocytes, which rely on laminin for anchoring through the integrin β1 protein.

### Inhibition of MMP2 reversed depigmentation in a novel mouse model of vitiligo

Finally, we ultilized K14-SCF transgenic mice, characterized by retained melanocytes in the epidermis, to investigate the impact of BM degradation on skin pigmentation. Murine Mock-Fbs and MMP2-Fbs were subcutaneously transplanted into the back skin of K14-SCF mice (**Fig 8a**). Subcutaneous transplantation of MMP2-Fbs in these mice resulted in vitiligo-like depigmentation (**Fig 8b**), with disrupted BM structure and absence of epidermal melanocytes (**Fig 8c**). Co-injection of MMP2-Fbs with MMP2 inhibitors (SB-3CT or tanomastat) effectively reversed the depigmentation (**Fig 8d, 8e and 8f**), supporting the crucial role of MMP2 in BM disruption and melanocyte loss. These findings establish the K14-SCF mouse model as a valuable tool for studying vitiligo pathogenesis and highlight MMP2 inhibition as a potential therapeutic approach for vitiligo treatment.

**Figure 8.**
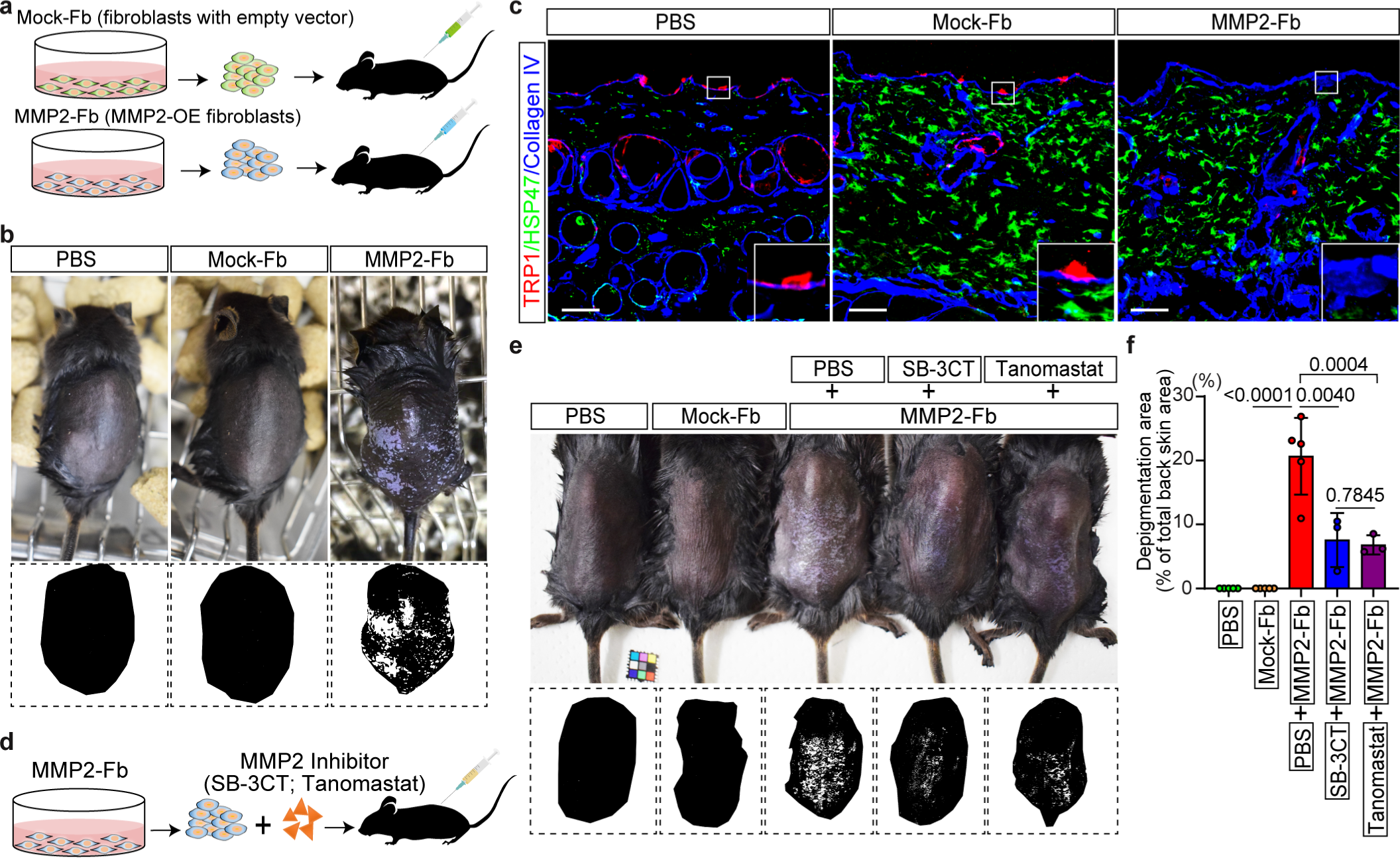
Subcutaneous inoculation of mice with MMP2-overexpressing fibroblasts induced a vitiligo phenotype that could be reversed by inhibition of MMP2 **a,** Schematic diagram of the subcutaneous fibroblast inoculation mouse model. **b,** Macro appearance of the mice at 3 weeks after fibroblast inoculation. The bottom panels show the corresponding threshold binarization images of the back skin of the mice. **c,** Representative immunofluorescence staining of TRP1, HSP47, and collagen IV in mouse skin sections. TRP1 is stained red, HSP47 is stained green, and collagen IV is stained blue. The area in the white rectangle is shown in the corresponding lower panel. Scale bars: 50 μm. **d,** Schematic of the experimental procedure to treat mice with MMP2 inhibitor. **e,** Macroscopic images of the mice at 2 weeks after subcutaneous inoculation. The bottom panels show the corresponding threshold binarization images of the back skin of the mice. **f,** Quantitation of the ratio of depigmentation area on the back skin of the mice. Data in **f** are shown as mean ± standard deviation.

## Discussion

The morphological abnormalities of the BM in vitiliginous skin have been previously reported, with thickening(*16, 17*), damage(*18*), or branching(*19*) observed. However, the underlying mechanisms of BM disruption and its role in the pathophysiology of vitiligo have remained unclear. In this study, we confirmed the abnormal BM structure in vitiliginous skin and identified that the disorganized BMZ prevents melanocytes from adhering to the BM via integrin β1–laminin binding. Furthermore, we discovered that the disorganized BM architecture is caused by increased MMP2 expression in dermal fibroblasts. Our findings shed light on the mechanisms underlying melanocyte disappearance in vitiligo and provide insights into potential therapeutic targets.

Previous studies have described melanocyte detachment in vitiligo, but they mostly focused on melanocyte–keratinocyte adhesion(*35, 36*). We addressed this gap by investigating melanocyte–BM adhesion and verified melanocyte detachment in vitiligo. Unlike previous studies that reported decreased expression of adhesion molecules or the melanocyte–BM adhesion molecule DDR1, our *in situ* PLA study revealed that the reduction in melanocyte adhesion was primarily due to the disorganization of the BMZ architecture. Our results suggest that the structural abnormality of the BM, rather than altered expression levels of adhesion molecules, is the primary cause of decreased melanocyte adhesion in vitiligo.

MMP2 and MMP9 are the main MMPs involved in breaking down BM components(*22, 37*). Our study found that MMP2 was dramatically elevated in the dermis of vitiliginous skin, and MMP9 expression was slightly increased as well, regardless of whether the vitiligo was stable or progressive. While previous studies have linked decreased E-cadherin expression in melanocytes(*36*) and increased MMP9 expression in vitiliginous skin(*38*) to melanocyte detachment, our findings demonstrate that the detachment of melanocytes is primarily related to their weak attachment to the disorganized BM, mediated by integrin β1–laminin binding and DDR1–collagen IV binding. This supports the notion that MMP2-induced BM disruption plays a crucial role in melanocyte disappearance in vitiligo.

The elevated expression of MMP2 in vitiliginous dermal fibroblasts highlights their potential contribution to vitiligo’s pathogenesis. Dermal fibroblasts play a crucial role in skin maintenance and healing, and aberrant behavior of these cells has been noted in vitiliginous skin(*39, 40*) (*41*). Our findings propose that increased MMP2 expression in dermal fibroblasts could disrupt the BM structure, resulting in melanocyte detachment. Cytokines like interleukin (IL)-1β(*42, 43*), IL-6(*42*), tumor necrosis factor (TNF)-α(*43, 44*), and TGF-β(*45, 46*) have been reported to stimulate MMP2 expression. Disruption of the BMZ architecture led to shifts in adhesive molecule expression in basal keratinocytes **(Fig 6)**. Likewise, heightened cytokine levels, including IL-1β, TGF-β1, TNF-α, and IL-6, were observed in vitiliginous skin lesions’ epidermis **(Extended Data Fig 1)**. Previous research aligns with our findings, suggesting disruption of the epidermal BM modulates interactions between epidermal keratinocytes and the extracellular matrix(*47*). Additionally, significant BM deterioration in pathological settings has been associated with increased cytokine release from epidermal cells(*47–49*). These cytokines might potentially spur MMP2 expression in dermal fibroblasts *via* paracrine regulation.

Hence, we present a proposed vicious cycle (**Fig 9**) where MMP2 instigates the BMZ, prompting changes in keratinocyte behavior. This alteration triggers cytokine release and upregulation of MMP2 in fibroblasts, perpetuating an unceasing cycle. This impaired BM repair gives rise to a pathological loop of cytokine and protease secretion, culminating in stable vitiligo, an obstinate skin condition. Local stress and microinflammation might incite this cycle. The NLRP3 and NLRP1 inflammasomes, activated by external triggers, initiate microinflammation and innate immunity, prompting increased cytokine levels like IL-1β, IL-6, and TNF-α in the epidermis(*50, 51*). These cytokines could potentially induce MMP2 expression in dermal fibroblasts, initiating the vitiligo development cycle. Further research is necessary to validate this hypothesis.

**Figure 9.**
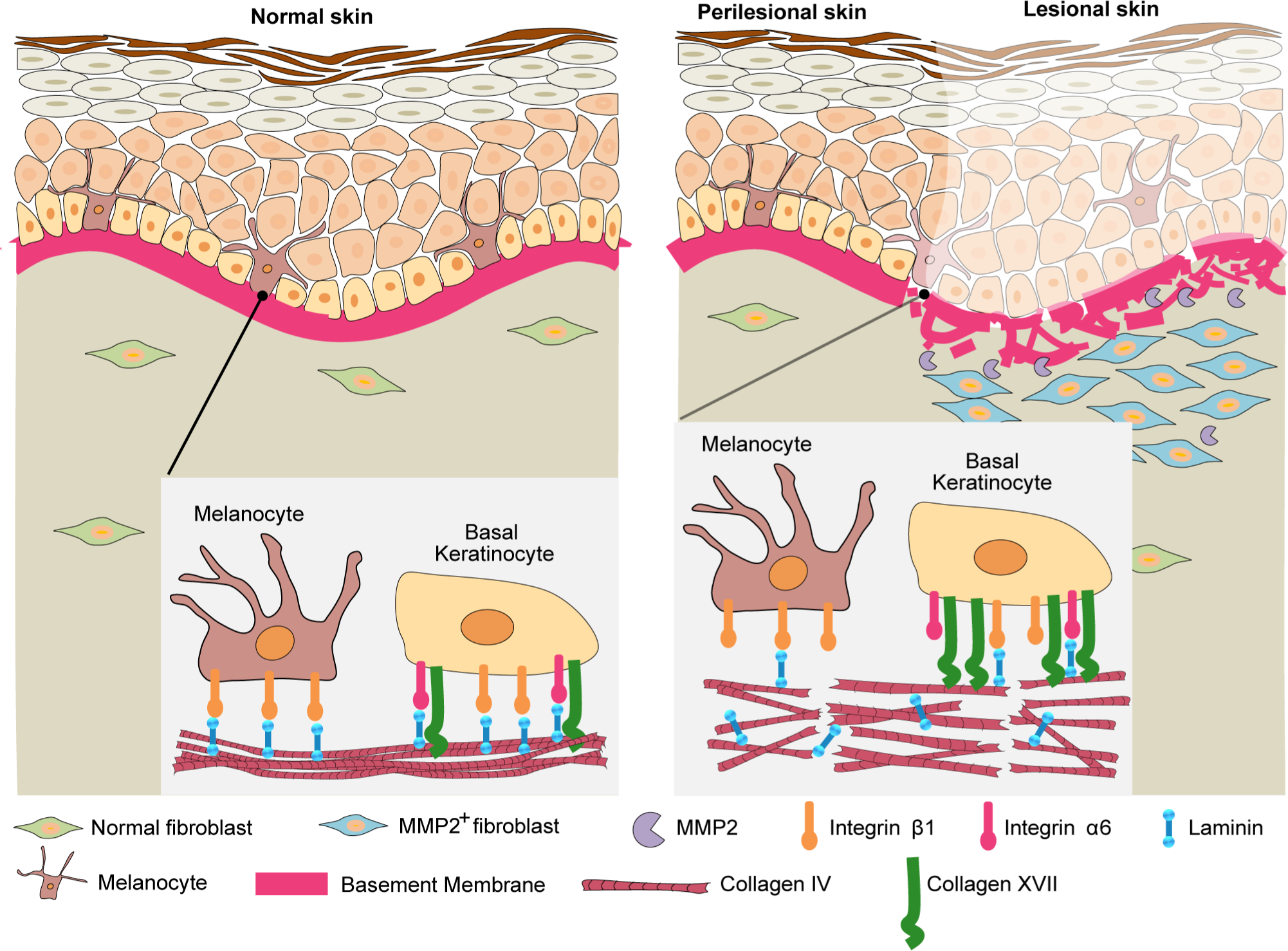
Summary illustration of MMP2-induced melanocyte disappearance in vitiligo. Schematic representation of normal skin and vitiliginous skin. Melanocytes are located at the basal layer of the epidermis interacting with the BM. In normal skin, laminin is normally distributed on collagen IV. Basal melanocytes attach to the BM mainly by integrin β1–laminin binding. In vitiliginous skin, collagen IV is degraded into fragments by the increased MMP2^+^ dermal fibroblasts, the laminin distribution is disrupted, and it is difficult for basal melanocytes to attach to the BM through integrin β1–laminin binding. Melanocytes detach from the BM and disappear from the epidermis. Although integrin β1–laminin binding and integrin α6–laminin binding are diminished in basal keratinocytes of vitiliginous skin, collagen XVII–collagen IV binding is enhanced as a compensatory mechanism.

Skin needling, laser ablation, and combined phototherapies with topical agents like steroids, tacrolimus, and 5-fluorouracil exhibit promising outcomes in stable vitiligo treatment(*52–54*). These methods stimulate the dermis, inducing biological changes in fibroblasts, the BM, and keratinocytes. This disruption in the pathogenic cycle aids BMZ repair. Inhibitors of inflammatory pathways, such as JAK inhibitors, may partially alleviate vitiligo. Restoring the BM appears pivotal for melanocyte repopulation. Targeting MMP2 expression and activation through pharmacologic and molecular interventions holds potential for vitiligo prevention and treatment. Normalizing MMP2 expression is crucial for a lasting cure. Clinical evaluation of MMP2 inhibitors shows promise, yet careful consideration of dosage and administration (e.g., topical application) is vital to minimize side effects and maintain melanocyte balance.

In our study, heightened MMP2 expression was evident in protein extracts from vitiligo lesions in comparison to healthy skin. Notably, the prevalent form of MMP2 in these extracts was the active 62- kDa variant, indicating localized activation of the latent (pro) form within the dermis of affected skin. The activation of pro-MMP2 is influenced by various factors, including TIMP-2, MT1-MMP, organomercurials, MMP7, and MMP3(*21*). Additional investigations are necessary to ascertain the potential involvement of these factors in MMP2 activation within vitiliginous skin.

In conclusion, our study provides valuable insights into the mechanisms of melanocyte disappearance in vitiligo. Disruption of the BM architecture, primarily caused by increased MMP2 expression in dermal fibroblasts, plays a crucial role in melanocyte detachment. These findings open new avenues for potential therapeutic strategies targeting MMP2 and BM repair to treat vitiligo. However, further research is needed to fully understand the complex interactions and mechanisms involved in vitiligo pathogenesis.

## Materials & Methods

### Human skin specimens

Histo-immunofluorescence staining was performed with frozen biopsy samples obtained from lesional and perilesional skin of well-defined patients with non-segmental vitiligo (n = 13, 7 in progressive state and 4 in stable state) (**Table 1**) and from corresponding sites on healthy donors (n = 5; 2 from the abdomen, 2 from an arm, and 1 from the chest). Three-millimeter punch fresh skin biopsy specimens of lesional skin from confirmed patients with vitiligo (n = 3) and samples from corresponding sites on healthy donors (n = 3) were used for protein extraction. One-millimeter punch skin biopsy specimens of lesional skin from confirmed patients with vitiligo (n = 5) and samples from corresponding sites on healthy donors (n = 5) were used for electron microscopy and immuno-electron microscopy analyses. The progressive state of vitiligo was defined by the development of new lesions or the extension of pre-existing lesions in the last 6 months, and the stable state was defined by no increase in the size of existing lesions and an absence of new lesions in the past 1 year. Written informed consent was obtained from all participants prior to study inclusion. The study was approved by the Medical Ethics Committee of Osaka Metropolitan University (No. 4152).

**Table 1.** Clinical characteristics of the patients.

| Patient No. | Sex | Age<br>(years) | Type | Biopsy site | Activity |
| --- | --- | --- | --- | --- | --- |
| 1<sup>§</sup> | M | 72 | Non-segmental vitiligo | Back | Progressive |
| 2<sup>§</sup> | F | 49 | Non-segmental vitiligo | Lumbar | Stable |
| 3* | M | 44 | Non-segmental vitiligo | Back | Progressive |
| 4* | F | 33 | Non-segmental vitiligo | Axilla | Progressive |
| 5* | F | 71 | Non-segmental vitiligo | Elbow | Stable |
| 6<sup>§</sup> | F | 23 | Non-segmental vitiligo | Abdomen | Stable |
| 7<sup>§</sup> | M | 30 | Non-segmental vitiligo | Lateral abdomen | Progressive |
| 8<sup>§</sup> | F | 20 | Non-segmental vitiligo | Abdomen | Progressive |
| 9 | M | 68 | Non-segmental vitiligo | Back | Progressive |
| 10 | F | 49 | Non-segmental vitiligo | Abdomen | Progressive |
| 11 | F | 62 | Non-segmental vitiligo | Neck | Stable |

\*Samples for full skin protein extract were obtained from these three patients.
<sup>§</sup>Samples for electron microscopy were from these five patients.

### Fluorescent Immunohistochemistry Staining

Human skin tissue and 3D skin samples embedded in OCT compound (Sakura Fineteck, Tokyo, Japan) were cut into 6-μm sections for immunohistochemistry analyses. After fixation with 4% paraformaldehyde (163-20145, Wako, Osaka, Japan) for 10 min and blocking with 5% BSA for 30 min at room temperature, these sections were incubated with 1:100 diluted primary antibodies specific for collagen IV (ab6586, Abcam, Cambridge, MA, USA), TRP1 (SIG-38150, Bio Legend, San Diego, CA, USA), MMP2 (ab86607, Abcam), MMP9 (ab119906, Abcam), HSP47 (ADI-SPA-470, Enzo Life Sciences, Farmingdale, NY, USA), CD31 (M0823, Dako, Santa Clara, CA, USA), laminin 332 (ab78286, Abcam), melan-A (SIG-38160, Bio Legend), cleaved-caspase 3 (#9664, Cell Signaling Technology, Danvers, MA, USA), DDR1 (PA5-29284, Thermo Fischer Scientific, Waltham, MA, USA), integrin β1 (ab30388, Abcam), Laminin (ab11575, Abcam), integrin α6 (ab20142, Abcam), and collagen XVII (HPA043673, Sigma-Aldrich, Burlington, MA, USA) and then with the appropriate secondary antibody (Alex Fluor 488/555/647 anti-rabbit IgG, anti-mouse IgG1, anti-mouse IgG2a, anti-mouse IgG2b, or anti-goat IgG; 1:1,000, Invitrogen, Grand Island, NY, USA). Nuclei were counterstained with DAPI (#62248, Thermo Fischer Scientific). Murine skin tissues embedded in OCT compound were cut into 6-μm sections for immunohistochemistry analyses. After fixation and blocking, these sections were incubated with 1:100 diluted primary antibodies specific for TRP1 (SIG- 38150, Bio Legend), HSP47 (ADI-SPA-470, Enzo Life Sciences), and collagen IV (ab19808, Abcam), and then with the appropriate secondary antibody (Alex Fluor 488 anti-mouse IgG2b, Alex Fluor 555 anti-mouse IgG2a, or Alex Fluor 647 anti-rabbit IgG; 1:1,000, Invitrogen). Slides were observed using an LSM 710 Zeiss confocal laser microscope. Images were captured using Zeiss ZEN 2012 software (Carl Zeiss Inc., Jena, Germany). Counting of positively stained cells and measurement of mean fluorescence intensity were carried out using Image J Software (NIH, http://rsb.info.nih.gov/ij/index.html).

### Electron Microscopy

Punch biopsy skin samples (1 mm) were fixed with 2.5% glutaraldehyde in 0.1 M phosphate-buffered saline (PBS) at pH 7.4 for 2 h and then washed overnight with PBS. Samples were then post-fixed in 1% OsO4 (157-01141, Wako) in PBS for 1h. After washing with DDW, samples were dehydrated with a graded ethanol series and embedded in EPON (TAAB EPON, 342-2, Nisshin EM, Tokyo, Japan). Ultrathin sections (70–80 nm) were cut and transferred to grids, dual-stained with uranyl acetate and lead citrate, and visualized on an FEI Talos F200C G2 Transmission Electron Microscope (Thermo Fisher Scientific). The width of lamina lucida was measured using Image J Software (NIH, http://rsb.info.nih.gov/ij/index.html).

### Pre-Embedding Immunoelectron Microscopy

Pre-embedding gold enhancement immunogold labeling was performed as detailed previously(*55*) with a slight modification. Briefly, skin tissues were fixed in 4% paraformaldehyde (163-20145, Wako) for 30 min at room temperature and embedded in OCT compound. Frozen sections (7 μm) were permeabilized in 0.25% saponin (47036, Sigma-Aldrich) for 30 min and then blocked for 30 min in PBS containing 0.1% saponin, 10% bovine serum albumin (013-25773, Wako), 10% normal goat serum (16210-072, Thermo Fisher Scientific), and 0.1% cold water fish skin gelatin (G7041, Sigma-Aldrich). Samples were incubated with anti-collagen IV (1:100; ab19808, Abcam) or anti-laminin (1:100; ab11575, Abcam) overnight at 4℃ in a blocking solution and then with colloidal gold-conjugated anti-rabbit IgG (1.4-nm diameter; 1:200; #2003, Nanoprobes, Yaphank, NY, USA) in a blocking solution for 2 h at room temperature. The signal was intensified with the Gold Enhancement EM Kit (#2113, Nanoprobes) for 4 min at room temperature. The specimens were post-fixed in 1% OsO4, dehydrated in a series of graded ethanol solutions, and embedded in EPON (TAAB EPON, 342-2, Nisshin EM). Ultrathin sections (70 nm) were collected and stained with uranyl acetate and lead citrate. Images were taken with an FEI Talos F200C G2 Transmission Electron Microscope (Thermo Fisher Scientific) equipped with an Advanced Microscopy Techniques charge-coupled device-based camera system. The quantitation analyses of positive gold particles were carried out using Image J Software (NIH, http://rsb.info.nih.gov/ij/index.html).

### Western Blot Analyses

For the preparation of protein samples, punch fresh skin biopsy specimens were extracted as described previously(*56*). Five micrograms of extracted protein was used for western blot analysis with primary antibody against MMP2 (1:200; ab92536, Abcam). The signal intensity of bands was quantified using the Image J densitometry software (NIH).

### Cell Culture and Establishment of MMP2-Overexpressing Cells

Normal human dermal fibroblasts (Neonatal, Primary, FC-0001, Lifeline Cell Technology, Carlsbad, CA, USA) were maintained in MEM (M4655, Sigma-Aldrich) with 1% NEAA (139-15651, Wako), 5% FBS (10270-106, Gibco, Thermo Fisher Scientific), 1 mM Sodium Pyruvate (11360-070, Gibco, Thermo Fisher Scientific), and 1% penicillin/streptomycin (168-23191, Wako). C57BL/6 mouse dermal fibroblasts (Neonatal, Primary, SCR-M230057, ScienCell Research Laboratories, Carlsbad, CA, USA) were maintained in Fibroblast Medium-2 (2331, ScienCell Research Laboratories). Cells were incubated at 37°C in a humidified incubator supplied with 5% CO2. Normal human dermal fibroblasts and C57BL/6 mouse dermal fibroblasts (2 × 10^6^ cells/well on 10- cm plates) were transfected with 20 μg human MMP2 expression or murine MMP2 expression episomal vector DNA or empty episomal vector DNA (customized pEBMulti-Puro, Fujifilm Wako Pure Chemical Corporation, Osaka, Japan) mixed with P3000 and Lipofectamine 3000 (Lipofectamine^TM^ 3000, Thermo Fisher Scientific). Successfully transfected cells were selected with 1 μg/ml puromycin according to the manufacturer’s instructions. The expression of the MMP2 proteins in cell lysate and culture medium were confirmed by western blot analyses using antibody against MMP2 (ab92536, Abcam).

### Reconstructed Three-Dimensional Human Full-Thickness Skin Equivalents

A reconstructed 3D human full-thickness skin equivalent model was generated as described previously(*57*) with modification. Collagen gels were prepared from an acid collagen solution (IAC- 50, Koken Ltd., Tokyo, Japan) according to the manufacturer’s protocol. Then, 2.5 × 10^5^ fibroblasts were cultured on collagen gel in MEM (M4655, Sigma-Aldrich) with 1% NEAA (139-15651, Wako), 5% FBS (10270-106, Gibco, Thermo Fisher Scientific), 1 mM sodium pyruvate (11360-070, Gibco, Thermo Fisher Scientific), and 1% penicillin/streptomycin (168-23191, Wako). This medium was changed every other day after fibroblast mounting. Two days after the dermal component was prepared, a cell suspension of 8.4 × 10^5^ keratinocytes and 1.2 × 10^5^ melanocytes in 60 μl medium composed of a 1:1 mixture of Epilife and Medium 254 was seeded onto the concave surface of the contracted gel. After 30 min, 2 ml culture medium was added to the inside and outside of the transwell, and the submerged culture was maintained for 2 days. The culture medium was composed of DMEM/F-12 (3:1 vol/vol; D5921, N4888, Sigma-Aldrich) containing 0.3% FCS, 0.1 mM ethanolamine, 0.1 mM *o*- phosphoethanolamine, 0.4 μg/ml hydrocortisone, 5 μg/ml insulin, 0.09 mM adenine, 5 μg/ml transferrin, 20 pM triiodothyronine, 0.64 mM choline chloride, 0.675 mM serine, 20 μg/ml linoleic acid/BSA, 50 μg/ml ascorbic acid, 7.25 mM glutamine, 10 nM progesterone, 100 I.U./ml penicillin, and 100 μg/ml streptomycin. The thus-formed early skin equivalents were then raised to the air-liquid interface and placed in cornification medium composed of a 1:1 mixture of Ham’s F-12 and DMEM supplemented with 0.3% FCS, 0.1 mM ethanolamine, 0.1 mM *o*-phosphoethanolamine, 0.4 μg/ml hydrocortisone, 5 μg/ml insulin, 0.09 mM adenine, 5 μg/ml transferrin, 20 pM triiodothyronine, 0.64 mM choline chloride, 0.75 mM serine, 20 μg/ml linoleic acid/BSA, 50 μg/ml ascorbic acid, 7.25 mM glutamine, 0.75 mM calcium chloride, 100 I.U./ml penicillin, and 100 μg/ml streptomycin. The medium was subsequently changed every other day. One week after the air-liquid interface incubation, the skin equivalents were harvested and subjected to macroscopic examination and further histology analyses. All skin equivalent experiments were performed in quintuplicate in five transwells.

To prepare the half-half 3D human skin equivalent, MMP2-overexpressing fibroblasts embedded in collagen gel were incubated at 37℃ for 30 min. The early-formed dermal tissue construct was then cut into half size and transferred to one side of a culture transwell insert, and freshly prepared collagen solution containing normal fibroblasts was injected into the other side of the transwell insert. After 2 days of culture, keratinocytes and melanocytes were seeded as described above.

### Ex Vivo Human Skin Explant Culture

Skin from the lateral abdomen was obtained with the informed consent of patients undergoing abdomen reduction surgery, in accordance with the ethical guidelines of the Research Ethics Committee of Osaka Metropolitan University (No. 4152). The explants were cut (1.0 cm^2^) and cultured at an air-liquid interface on culture inserts (filter pore size 8 μm; #141082, Thermo Fisher Scientific) with a 1:1 mixture of Ham’s F-12 and DMEM supplemented with 0.3% FCS, 0.1 mM ethanolamine, 0.1 mM *o*-phosphoethanolamine, 0.4 μg/ml hydrocortisone, 5 μg/ml insulin, 0.09 mM adenine, 5 μg/ml transferrin, 20 pM triiodothyronine, 0.64 mM choline chloride, 0.75 mM serine, 20 μg/ml linoleic acid/BSA, 50 μg/ml ascorbic acid, 7.25 mM glutamine, 0.75 mM calcium chloride, 100 I.U./ml penicillin, and 100 μg/ml streptomycin. The insert was set on a 12-well plate (3401, Corning Inc., Corning, NY, USA) and cultured at 37℃ in a humidified incubator with 5% CO2 for 7 days.

For fibroblast inoculation, normal fibroblasts and MMP2-overexpressing fibroblasts were labeled with the red fluorescent dye PKH26 (MINI26-1KT, Sigma-Aldrich) according to the manufacturer’s instructions and then subcutaneously injected into cultured human skin explants at a density of 5 × 10^6^ cells/ml in 100 μl using disposable 35G needles (ReactSystem, Osaka, Japan).

### *In Situ* Proximity Ligation Assay

Multicolor *in situ* PLA was performed using the Duolink PLA Multicolor detection kit (Duolink^®^ PLA^®^ Multicolor Reagent Pack, DUO96000, Sigma-Aldrich) as described previously(*58*) with slight modification. Briefly, frozen skin sections were blocked with blocking solution (DUO82007, Duolink, Sigma-Aldrich) for 30 min at 37 °C and then incubated overnight at 4°C with primary antibodies pre-conjugated with Red Oligo A (DUO86010A, Sigma-Aldrich) or Red Oligo B (DUO86010B, Sigma-Aldrich). After washing with Duolink *In Situ* Wash Buffer A (DUO82046, Sigma-Aldrich), ligation and amplification reactions were performed using a Duolink PLA Multicolor detection kit (DUO96000, Sigma-Aldrich).

For co-staining with melanocyte marker after amplification and washing, sections were incubated with the Alexa Fluor 488-conjugated antibody cocktail (MART1 + Tyrosinase + gp100) [Alexa Fluor^®^ 488] (1:100 dilution; NBP2-34681AF488, Novus Biologics, San Jose, CA, USA) for 2 h. After 30 min of washing with Duolink *In Situ* Wash Buffer B (DUO82048, Sigma-Aldrich), sections were mounted using Duolink *In Situ* Mounting Medium with DAPI (DUO82040, Sigma-Aldrich). Then, slides were observed using an LSM 710 Zeiss confocal laser microscope. Images were captured using Zeiss ZEN 2012 software (Carl Zeiss Inc., Jena, Germany). Mean fluorescence intensity measurements were carried out using Image J Software (NIH, http://rsb.info.nih.gov/ij/index.html).

### Animals and *In Vivo* Experimental Procedures

Krt14-Kitl mice, a C57BL/6-background congenic strain that expresses membrane-bound stem cell factor under the keratin14 promoter to retain epidermal melanocytes and thus have pigmented skin, were purchased from the RIKEN BioResource Research Center (No. RBRC00694, RIKEN BRC, Tsukuba, Ibaraki, Japan). PBS, Mock-Fbs, or MMP2-Fbs (5 × 10^6^ cells/ml in 100 μl PBS) was subcutaneously injected into the back skin of male 7-week-old Krt14-Kitl mice.

For treatment, 5 μM SB-3CT dissolved in DMSO (HY-12354, MedChemExpress, Monmouth junction, NJ, USA) or 5 μM Tanomastat dissolved in DMSO (T006900, Toronto Research Chemicals, Toronto, Canada) was subcutaneously co-injected with Mock-Fbs or MMP2-Fbs (5 × 10^6^/ml in 100 μl PBS) into the back skin of Krt14-Kitl mice. Two weeks after these treatments, the back skin of the mice was depilated and photographed, and the depigmentation area (n = 5 mice per group) was measured using binarization and different global thresholding values (Image J Software, NIH, http://rsb.info.nih.gov/ij/index.html). Experiments were conducted in accordance with the Guiding Principles for the Care and Use of Laboratory Animals, and the experimental protocol used in this study was approved by the Committee for Animal Experiments at Osaka Metropolitan University (Permit No. 18042, Osaka, Japan).

### Statistical Analysis

Experiments were repeated at least three times. Data are presented as the mean ± standard deviation. One-way analysis of variance with Dunnet’s test or unpaired Student’s t-test (two-tailed) was used for statistical analyses. A value of p < 0.05 indicated statistical significance. All statistical analyses for this study were performed using GraphPad Prism version 8.0.0 for Windows (GraphPad Software Inc., San Diego, CA, USA).

## Supplementary Information

### Materials & Methods

#### Fluorescent Immunohistochemistry Staining

Human skin tissue embedded in OCT compound (Sakura Fineteck, Tokyo, Japan) was cut into 6- μm sections for immunohistochemistry analyses. After fixation with 4% paraformaldehyde (163- 20145, Wako, Osaka, Japan) for 10 min and blocking with 5% BSA for 30 min at room temperature, these sections were incubated with 1:100 diluted primary antibodies specific for TGF-β1 (ARG66427, Arigobio, Hsinchu, Taiwan), TNF-α (60291-1-Ig, Proteintech, Rosemont, USA), IL-1β (16806-1-AP, Proteintech), and IL-6 (21865-1-AP, Proteintech) and then incubated with the appropriate secondary antibody (Alex Fluor 488/555 anti-rabbit IgG, anti-mouse IgG1, anti-mouse IgG2a; 1:1,000, Invitrogen, Grand Island, NY, USA). Nuclei were counterstained with DAPI (#62248, Thermo Fischer Scientific). Slides were observed using an LSM 710 Zeiss confocal laser microscope. Images were captured using Zeiss ZEN 2012 software (Carl Zeiss Inc., Jena, Germany). Counting of positively stained cells and measurement of mean fluorescence intensity were carried out using Image J Software (NIH, http://rsb.info.nih.gov/ij/index.html).

---

**Extended Data Fig. 1.**
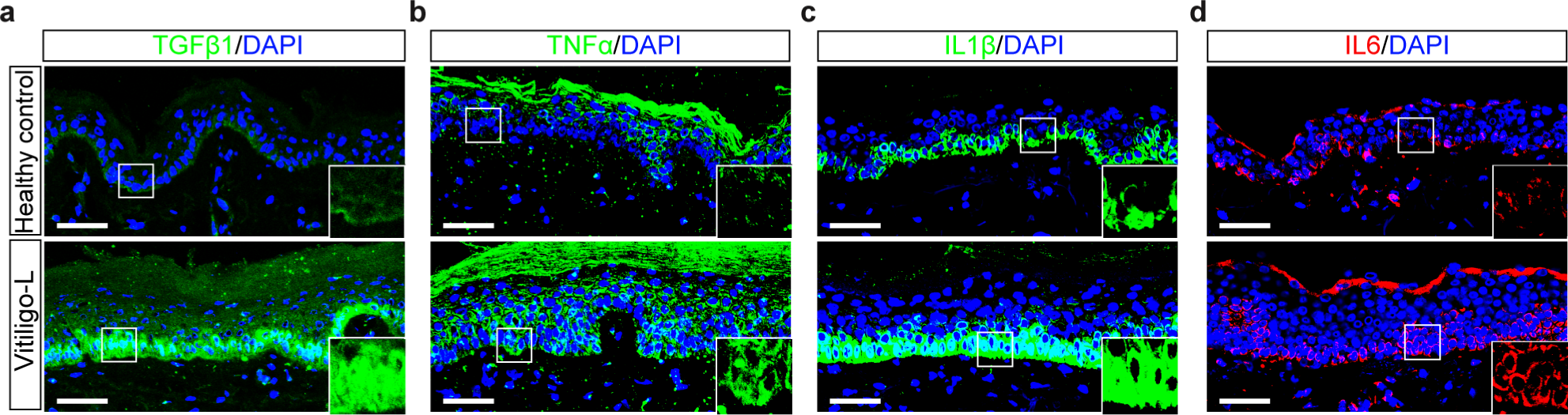
Increased cytokine expression in the epidermis of vitiliginous skin. Representative immunofluorescence staining of TGF-β1 (**a**), TNF-α (**b**), IL-1β (**c**), and IL-6 (**d**) in skin sections from healthy controls and lesional skin sections from patients with vitiligo. Nuclei were counterstained in blue with DAPI. The area in the white rectangle is shown on the lower right. Scale bars in **a, b, c**, and **d**: 50 μm.

## Notes

### Competing Interest Statement

The authors have declared no competing interest.

### Summary of Updates

Abstract, Introduction, Discussion revised

